# ACHT2 deactivates carbon assimilation during light-dark transition

**DOI:** 10.64898/2026.09.02.748655

**Authors:** Moshe Steinberg, Matanel Hipsch, Misgav Keisar, Nardy Lampl, Shilo Rosenwasser

## Abstract

In chloroplasts, photosynthetic efficiency relies on a delicate balance between reductive and oxidative thiol-based signaling networks. Members of the high-midpoint-potential atypical thioredoxins (Trxs) were shown to oxidize photosynthetic enzymes by channeling reducing equivalents to H_2_O_2_ through 2-Cys-Prx activity. However, it remains unclear which atypical-Trx isoforms regulate Calvin-Benson cycle (CBC) inactivation, and whether they possess distinct functional specificities or operate redundantly in vivo. To resolve this, electron transport and carbon assimilation were continuously monitored during physiological dynamic light transitions in CRISPR-generated single, double and triple atypical Trx mutants. Notably, ACHT2 was identified as a primary determinant of CBC inactivation during dark-to-light transitions, as evidenced by the alleviation of redox-mediated bottlenecks in electron flow downstream of Fd and a lower CBC inactivation state in *acht2* plants, resembling the phenotype observed in plants lacking 2-Cys Prxs A and B (*2cpab*). In contrast, plants mutated in ACHT1, ACHT4, or TrxL2 displayed CBC inactivation kinetics comparable to those of the wild type. Furthermore, mutation of ACHT2 did not compromise plant fitness. In contrast, growth retardation was observed in the *acht1/acht4* double mutant, suggesting that the severe phenotype of *2cpab* does not arise from impaired CBC inactivation, but rather from disruption of oxidative regulation of other metabolic pathways mediated by distinct atypical Trxs. Collectively, these findings reveal a high degree of regulatory specificity within the chloroplast oxidative network and provide a foundation for a deeper understanding of how activation-inactivation cycles contribute to plant adaptation to dynamic light environments.

## Introduction

Plants in natural environments experience highly dynamic light conditions, necessitating frequent and rapid adjustments of chloroplast metabolism to optimize photosynthetic efficiency and prevent oxidative damage. At the core of this metabolic flexibility lies thiol-based redox regulation, a post-translational mechanism that synchronizes the activity of metabolic enzymes with the functional state of the photosynthetic electron transport chain (PETC) (Buchanan et al., 2016; Schürmann and Buchanan, 2008; Yoshida et al., 2019; Dietz and Pfannschmidt, 2011; Foyer and Noctor, 2009). This regulatory network operates by reversibly modifying the redox states of specific cysteine residues, thereby acting as a molecular switch that modulates chloroplast enzyme activity to balance carbon assimilation with energy dissipation (Wittenberg and Danon, 2008; Cejudo et al., 2021).

Upon illumination, reducing power generated by the PETC is channeled from ferredoxin (Fd) to thioredoxins (Trxs) via Fd-Trx reductase (FTR), as well as through the ferredoxin-NADP(+) reductase (FNR) and NADPH-dependent Trx reductase C (NTRC) pathway (Buchanan and Balmer, 2005, Serrato et al., 2004). This cascade orchestrates the rapid activation of essential Calvin-Benson cycle (CBC) enzymes, such as glyceraldehyde 3-phosphate dehydrogenase (GAPDH), fructose 1,6-bisphosphatase (FBPase), sedoheptulose 1,7-bisphosphatase (SBPase) and phosphoribulokinase (PRK), by reducing intramolecular disulfide bonds, essentially turning on CO_2_ assimilation in a light-dependent manner (Michelet et al. 2013; Gurrieri et al. 2021). Efficient deactivation of these pathways is necessary to prevent the futile depletion of metabolic intermediates when energy supply is interrupted. Yet, while the mechanisms of reductive activation are well established, the processes governing oxidative deactivation during light-to-dark transitions are less understood.

Recent works have demonstrated that the oxidation of redox-regulated proteins is catalyzed by a distinct oxidative network involving 2-Cys peroxiredoxins (Prx) and atypical Trx-like proteins (Ojeda et al., 2018; Eliyahu et al., 2015; Dangoor et al., 2012; Yoshida et al., 2018; Vaseghi et al., 2018). Indeed, a recent redox proteomic mapping highlighted the central role of 2-Cys Prxs in shaping this dynamic redox landscape, demonstrating their role in mediating oxidative signaling during light-to-dark transitions (Doron et al., 2025). 2-Cys Prxs utilize hydrogen peroxide (H₂O₂) as a terminal electron acceptor to continuously counterbalance reductive signals and act as a metabolic brake on carbon assimilation (Lampl et al., 2022). This reliance on H₂O₂ directly links metabolic regulation to chloroplastic reactive oxygen species (ROS) dynamics. Although historically viewed as incidental, detrimental by-products, ROS generated via mechanisms such as the water-water cycle (WWC) are increasingly recognized as vital signaling molecules (Mittler, 2017; Wrzaczek et al., 2013; Noctor et al., 2018; Hipsch et al, 2026). The physiological prominence of this oxidative system is highlighted in mutant lines lacking 2-Cys Prxs, which exhibit growth retardation, lower chlorophyll content, and altered photosynthetic induction kinetics (Pérez-Ruiz et al., 2017; Awad et al., 2015; Ojeda et al., 2018; Lampl et al., 2022). However, dark-induced oxidation of CBC enzymes was delayed but not completely abolished in 2-Cys Prxs A and B double mutants (*2cpab*) (Ojeda et al., 2018; Yoshida et al., 2018; Doron et al., 2025), suggesting that compensatory alternative pathways, possibly involving other plastid peroxidases or glutathione (GSH), can function when 2-Cys Prxs are unavailable.

Among the atypical Trxs are the atypical cysteine histidine-rich Trxs (ACHT) and TRX-like 2 (TRXL2) families, which are distinguished by non-canonical CXXC active sites and less negative redox potentials (Dangoor et al., 2009; Chibani et al., 2021; Yoshida et al., 2018). These atypical isoforms drive the oxidative network by preferentially transferring electrons from plastid proteins to 2-Cys Prxs (Ojeda et al., 2018; Eliyahu et al., 2015; Dangoor et al., 2012; Yoshida et al., 2018; Vaseghi et al., 2018).

Studies investigating the activity of atypical Trxs have yielded a complex picture of their target specificity. On the one hand, several isoforms appear to exert specific oxidative control. For example, thioredoxin-like2.1 (TrxL2.1) acts as a dedicated oxidizing factor for the γ-subunit of ATP synthase (CF1-γ) (Yoshida et al., 2018; Yokochi et al., 2021; Sekiguchi et al., 2022). Similarly, members of the atypical Cys-His-rich thioredoxin (ACHT) family have been assigned distinct targets; ACHT4 drives the oxidation of ADP-glucose pyrophosphorylase (AGPase) to attenuate starch synthesis (Eliyahu et al., 2015), while ACHT1 and ACHT2 target fructose 1,6-bisphosphatase (FBPase) (Yokochi et al., 2019; Yokochi et al., 2021). On the other hand, emerging evidence suggests a significant degree of functional redundancy. For example, recent analyses indicated that multiple isoforms, including ACHT3, ACHT4, and TrxL2.2, act redundantly to mediate dark-dependent oxidation through 2-Cys Prxs (Jiménez-López et al., 2025).

The current work utilized continuous, real-time physiological monitoring during dark-to-light transitions to capture the oxidative deactivation kinetics of CBC enzymes. By comparing wild-type *Arabidopsis thaliana* with CRISPR/Cas9-generated knockout mutant lines, ACHT2 was shown to be the primary inhibitor of CBC enzymes.

## Results

### Generation of *A. thaliana* ACHT and TRX-L CRISPR/Cas9 mutants

To investigate the physiological role of atypical Trxs in regulating photosynthesis-related reactions, a set of mutants was generated using a CRISPR/Cas9-mediated genome-editing approach (Ran et al., 2013). Specifically, ACHT1, ACHT2, ACHT4, TRXL2.1 and TRXL2.2 were targeted as large-scale transcriptional data indicated their high expression levels in leaves (Supp. Fig. 1). At least two independent homozygous knockout lines were isolated for each gene target, following Agrobacterium-mediated transformation and subsequent screening of T2 segregants, except for TRXL2.1, for which a single line was generated (Supp. Fig.2). In addition, to explore the possible functional redundancy between these genes, *acht1*/*acht2* and *acht1*/*acht4* double mutants (Supp. Fig.2), as well as the *acht1*/*acht2*/*acht4* triple mutant (Fig.1a) were generated. In all lines, the Cas9 cassette was segregated out based on mCherry fluorescence, resulting in null segregants. The complete list of lines used, including the specific mutation types and their corresponding positions, is provided in Supp.Table 1. In addition to these lines, the *2cpab* mutant line, which is mutated in 2-Cys Prx A and 2-Cys Prx B (Ojeda et al., 2018), was included in all experiments to assess the consequences of complete inhibition of the oxidative pathway.

**Figure 1:**
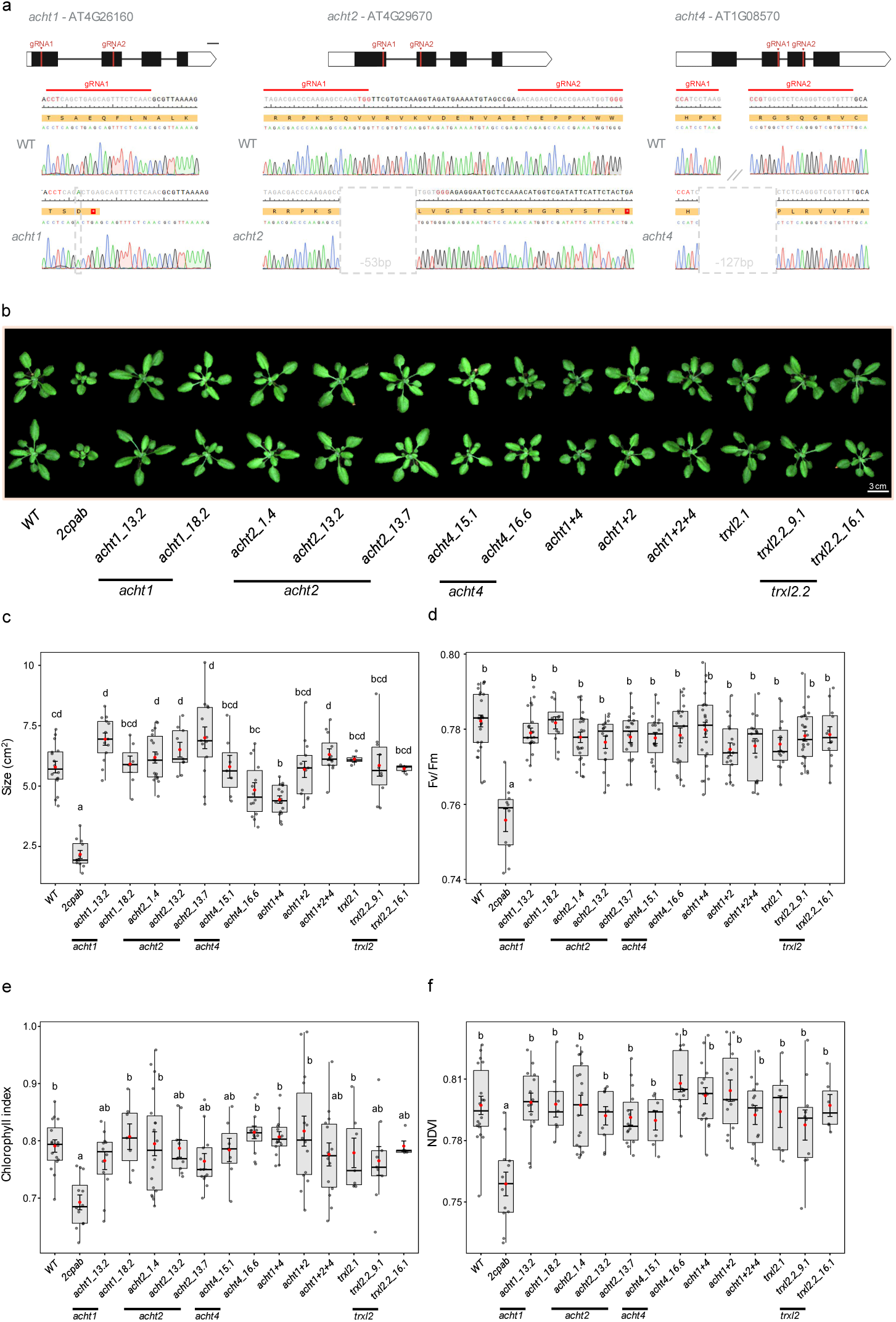
Phenotypic and physiological characterizationof acht and Trx-l2 mutants. (a) Sanger sequencing chromatograms confirmingthe targeted *acht1/acht2/acht4* mutations in the triple mutant line. (b) Representative visible phenotypes of WT, *2cpab* and mutant lines at 2 weeks post-germination. (c-f) Quantification of growth and physiological parameters across the mutant lines: (c) rosette size, (d) maximum quantum yield of photosystem II (Fv/Fm), (e) chlorophyll index and (f) normalized difference vegetation index (NDVI). Data are presented as box-and-whisker plots, which indicate the median and interquartile range, with individual biological replicates overlaid as grey dots (n = 5-20). Red dots denote the mean ± SE. Distinct letters indicate significant differences between lines (one-way ANOVAfollowed by Tukey’s HSD test; p < 0.05).

**Figure 2:**
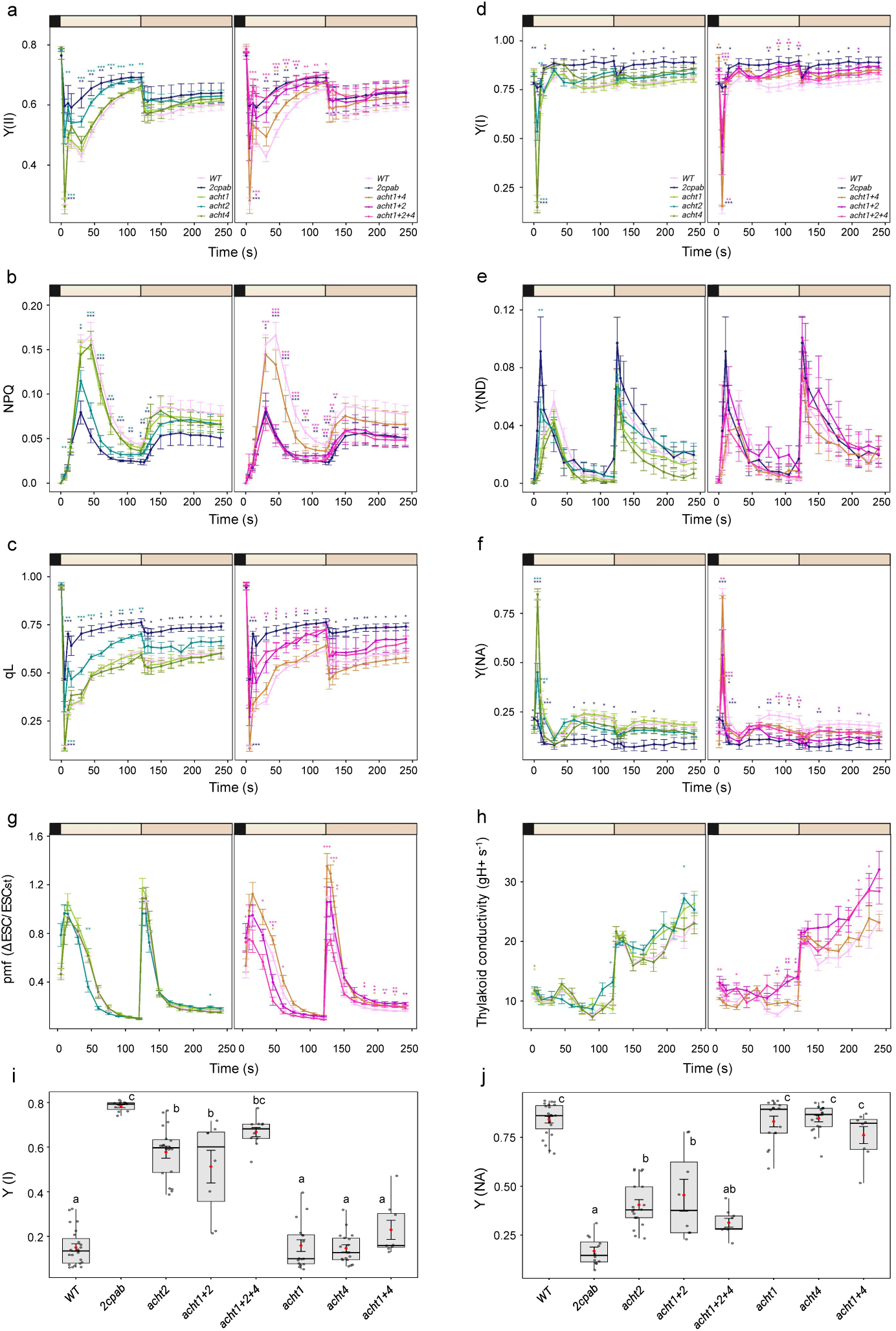
Photosynthetic electron transport and thylakoid proton dynamics under dark to low light transition. Analysis of kinetic photosynthesis parameters across the *acht* mutant lines under a gradient of actinic light intensities (0, 35, 80 µmol photons m−2 s−1). Panels display (a) quantum yield of photosystem II (PSII) (Y(II)), (b) non-photochemical quenching (NPQ), (c) fraction of open PSII reaction centers (qL), (d) quantum yield of photosystem I (PSI) (Y(I)), (e) PSI donor-side limitation (Y(ND)), (f) PSI acceptor-side limitation (Y(NA)), (g) proton motive force (pmf) and (h) thylakoid proton conductance (gH+). Colored bars above the graphs denote the applied actinic light intensities. Expanded view of (i) Y(I) and (j) Y(NA) during the first 5 s of illumination. Data are presented as the mean ± SE(n = 5-20 biological replicates). Statistical significance was assessed for panels (a-h) was assessed using Welch’s *t*-test with Bonferroni correction for multiple comparisons against the wild type and is marked as p < 0.05 (*), p < 0.01 (**), p < 0.001 (***). For panels (i-j), distinct letters indicate significant differences between all plant lines (one-way ANOVAfollowed by Tukey’s HSD test; *P* < 0.05). For the comprehensive response across all increasing light intensities, see Supplementary Figure 3.

Marked growth inhibition of over 60% compared with WT was measured in *2cpab* (Tukey’s HSD test, p<0.05, Fig1.b and c). *acht1/acht4* displayed an intermediate but significant growth reduction of 24%, while the remaining mutant lines showed no statistically significant differences from the WT (Fig1.b and c). Fv/Fm values across most lines remained within the normal physiological range of 0.77 to 0.8, while the *2cpab* mutant showed a slight reduction, with an average of 0.75 (Fig. 1d). Similarly, analysis of the red edge chlorophyll index and Normalized Difference Vegetation Index (NDVI) confirmed high vegetative vigor among the mutant lines, with values falling within the normal ranges of 0.65-0.9 and 0.70-0.82, respectively. However, *2cpab* displayed significantly lower, yet within-range, values across both indices compared to the WT (Fig. 1e-f). Overall, these results demonstrate highly competent photosynthetic tissues in the generated tested mutant lines and no severe phenotypes, indicating that essential cellular processes were not disrupted in the absence of atypical Trxs.

### ACHT2 regulates acceptor-side limitations under dark-to-light transition

Simultaneous measurement of chlorophyll fluorescence, P700 redox changes and electrochromic shift (ECS) absorbance during dark-to-light transitions and under incremental increases of light intensity were carried out to explore the role of atypical Trxs in regulating photosynthetic activity (Fig. 2 and Supp. Fig. 3 and 4). Notably, *acht4* displayed significantly lower PSI donor-side limitation Y(ND) than the WT under high light intensity (1200 µmol photons m⁻² s⁻¹), despite showing no alterations in proton motive force (pmf) or proton conductivity of the thylakoid membrane (gH⁺) (Supp. Fig. 3). These results may indicate reduced photosynthetic control (PCON) in *acht4*, independent of changes in the ΔpH gradient. Notably, this unique Y(ND) phenotype was absent in the *acht1*/*acht4* double mutant (Supp. Fig. 3), indicating compensatory dynamics between the specific isoforms.

**Figure 3:**
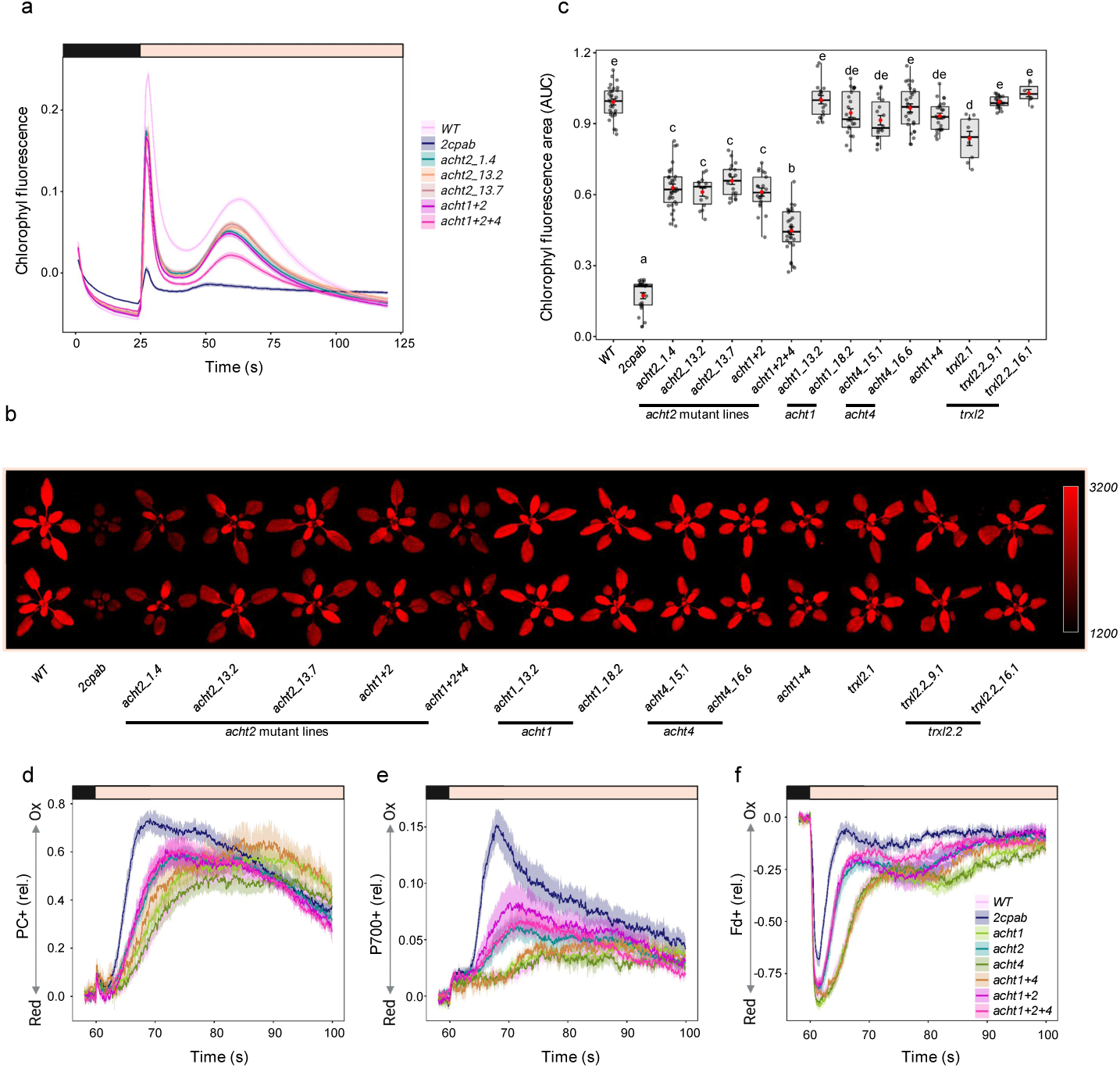
Induction kinetics of chlorophyll fluorescence, and redox state of Pc, p700 and Fd duringdark to low light transition. (a) Chlorophyll fluorescence induction curves (Kautsky effect) of dark-adapted *acht2* mutant lines transitioned to low light (35 µmol photons m−2 s−1). (b) Area under the fluorescence induction curve (25 s-100 s), serving as a quantitative proxy for the Kautsky effect. (c) False-color images of plants captured 5 s after the onset of actinic illumination. Pixel intensities reflect the transient quenching state mapped to a red color scale ranging from 1200 to 3200 arbitrary units. (d) Fraction of oxidized plastocyanin (Pc). (e) Fraction of oxidized p700. (f) Fraction of reduced ferredoxin (Fd). Data are presented as the mean ± SE(n = 8-32 biological replicates). Distinct letters indicate significant differences between lines (one-way ANOVAfollowed by Tukey’s HSD test; P < 0.05).

**Figure 4:**
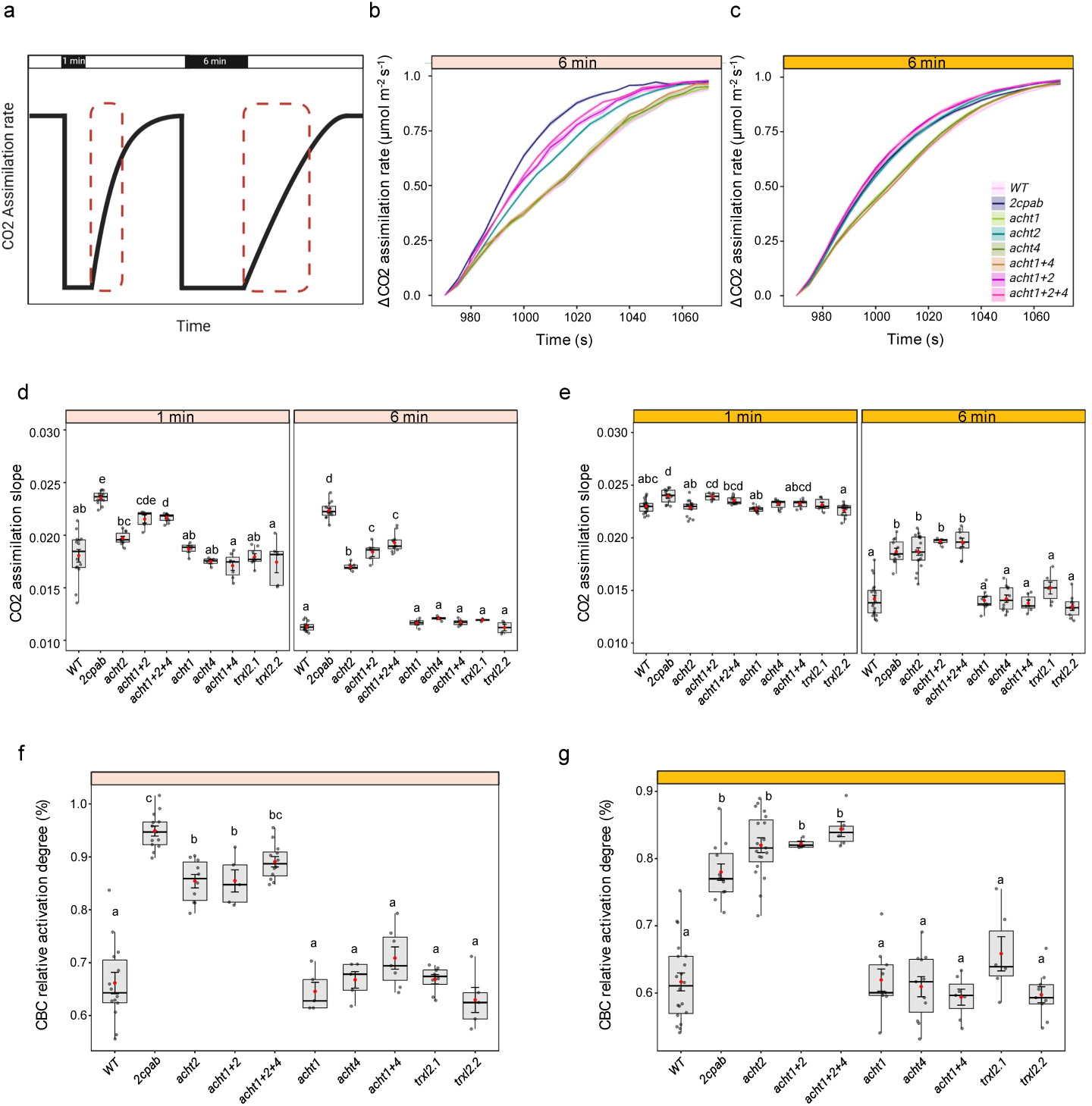
Loss of *acht2* delays the deactivation of carbon assimilation in darkness. (a) Schematic presentation of the experimental protocol. Red dashed rectangles denote the initial linear phase of CO• assimilation induction, from which the recovery slope was calculated following each dark interval. (b, c) Initial induction rates of CO• assimilation across the *acht* mutant lines after 6 min of dark adaptation, measured under subsequent (b) low (40 µmol photons m^⁻²^ s^⁻¹^) or (c) moderate (140 µmol photons m^⁻²^ s^⁻¹^) light intensities. To evaluate relative induction kinetics, all data were normalized to the final steady-state assimilation rate achieved under each respective light condition. (d, e) Comparison of CO• assimilation induction slopes following 1 min and 6 min of dark adaptation under (d) low and (e) moderate light intensities. (f, g) CBC relative activation degree under (f) low and (g) moderate light intensities. Data are presented as box-and-whisker plots, which indicate the median and interquartile range, with individual biological replicates overlaid as grey dots (n = 4 - 19). Red dots denote the mean ± SE. Distinct letters indicate significant differences between lines (one-way ANOVA followed by Tukey’s HSD test; P < 0.05).

Interestingly, during the dark-to-light transition, *2cpab* and *acht2* mutant lines exhibited higher photosystem II (PSII) quantum yield (Y(II)) than WT, which correlated with reduced NPQ induction (Fig. 2a-b). This divergence was even more pronounced for the fraction of open PSII centers (qL), indicating higher Qa oxidation states in these mutants during the transition to light (Fig. 2c). Furthermore, in WT, a transient spike in PSI acceptor-side limitation Y(NA) was observed, reflecting a temporary electron bottleneck at Fd, which subsequently relaxed. This spike was significantly attenuated in the *acht2* mutant lines and completely abolished in *2cpab* (Fig. 2f and j). This acceptor-side limitation (Y(NA))-dependent divergence was most prominent under low light conditions (<80 µmol photons m⁻² s⁻¹) and became less pronounced during the transition to higher light intensities (Supp. Fig.3).

In WT plants exposed to low light intensities, each stepwise increase in light triggered a transient pmf spike followed by rapid relaxation (Fig. 2g and h). *acht2* plants displayed a slightly reduced spike and faster relaxation than the WT, while the most pronounced *pmf* phenotype was observed in *acht1/acht2/acht4*, which maintained a significantly attenuated pmf under low light conditions (Fig. 2g). This pmf reduction was driven by an increase in gH⁺ (Fig. 2h), indicating elevated proton conductance. Notably, the *trxl2* mutant lines exhibited no significant differences compared to the WT, with the exception of slightly lower but significant Y(NA) values transiently observed upon dark-to-light induction. (Supp. Fig 4). Overall, Y(NA) was alleviated under low light in *acht2* plants, as reflected by a higher Qa oxidation state, faster pmf relaxation and consequently lower NPQ levels.

### Loss of ACHT2 alleviates redox-mediated bottlenecks in electron flow downstream of Fd

The relief of the Y(NA) blockage in the *acht2* mutant lines suggests that ACHT2 dictates the pace of acceptor-side activation. To further examine whether the electron transport chain is more poised to accept electrons in the absence of ACHT2, the transient rise and subsequent quenching of chlorophyll fluorescence (Kautsky et al., 1931), along with redox changes in plastocyanin (Pc), P700 and Fd at photosynthesis onset were monitored. As previously shown, the Kautsky effect was abolished in *2cpab* plants, suggesting sustained photochemical or non-photochemical quenching in the dark (Vaseghi et al., 2018), and indicating that disruption of redox regulation is manifested in altered chlorophyll fluorescence kinetics. While most of the single mutants displayed Kautsky induction kinetics indistinguishable from the WT (Supp. Fig. 5 a-d), *acht2* mutants displayed an intermediate pattern between *2cpab* and WT (Fig. 3a). Chlorophyll fluorescence induction dynamics, quantified by integration of the area under the chlorophyll fluorescence curve, was significantly reduced to 18% and 60% of the WT in *2cpab* and *acht2* lines, respectively. The *acht1/acht2* double mutant showed a pattern resembling that of the *acht2* mutant, while fluorescence quenching in the *acht1/acht2/acht4* triple mutant dropped to 45% of the WT signal, indicating partial degree of hierarchical redundancy among the ACHT genes. Alternatively, the lack of an effect in the single-mutant *acht1* and *acht4* lines (Fig. 3a and Supp. Fig. 5 a-d) may suggest that ACHT1 and/or ACHT4 regulate additional electron acceptors, a regulatory role that becomes more prominent in the *acht2* mutant background. Aside from a modest reduction to 84% of the WT signal in *trxl2.1*, no other lines showed significant differences from the WT (Fig. 3b). Because chlorophyll fluorescence under low-light conditions, where antenna light harvesting is not yet saturated, may indicate the recruitment of downstream electron sinks, these findings suggest that ACHT2 is a major redox regulator that maintains the PETC in a state poised to accept electrons.

Accordingly, the role of atypical Trxs in shaping electron fluxes was further investigated by measuring the redox states of Pc, P700 and Fd. In WT plants, a distinct lag in Fd oxidation was observed upon illumination, indicating the closed electron gate imposed by acceptor-side limitations (Fig. 3f). Both the 2cpab and acht2 mutant lines exhibited markedly accelerated Fd oxidation kinetics compared to WT, indicating a rapid alleviation of downstream acceptor-side limitations. The observed kinetics of Fd oxidation, which closely mirrored fluorescence quenching across all lines (Supp. Fig. 6), were in agreement with previous observations (Schreiber et al., 2017), demonstrating a strong correlation between downstream acceptor activation and chlorophyll fluorescence quenching. As for *trxl2*, the Pc, P700, and Fd redox kinetics patterns were similar to those of the WT (Supp. Fig. 7). No P700 oxidation was observed in WT upon transition to low light, suggesting that during the initial induction phase, the rate of its re-reduction by PSII equaled rate of downstream electron consumption. By comparison, *2cpab* and *acht2* lines exhibited a transient oxidation peak, indicating greater electron consumption than WT, with *acht2* showing a less pronounced peak than *2cpab* (Fig. 3e). Similar patterns were obtained for Pc oxidation (Fig. 3d). Overall, these observations suggest that the redox regulatory system imposes constraints on electron flow downstream of Fd, and that these constraints are alleviated in the absence of 2-Cys Prx and ACHT2, but not upon loss of other members of the atypical Trx family.

### ACHT2 is the primary oxidative inhibitor of the Calvin–Benson cycle (CBC)

The suggested relief of limitations in electron flow during photosynthetic induction in *acht2* plants, reflected in the reduced Y(NA), chlorophyll fluorescence signal and altered oxidation dynamics of Pc, P700 and Fd, prompted an examination of the deactivation dynamics of the CBC, the primary sink for photosynthetic reducing power. This was achieved by comparing CO_2_ assimilation rates in plants reilluminated after 1 and 6 min of dark adaptation, thereby capturing the activated and oxidatively inactivated states of CBC enzymes, respectively (Fig. 4a; Hipsch et al., 2026). When plants were reilluminated under low light intensity (40 µmol m⁻² s⁻¹), where photosynthetic electron flux is limited, WT-normalized CO_2_ assimilation slopes declined from 0.018 after 1 min of dark adaptation to 0.011 after 6 min (Fig. 4b and d), indicating a 34% reduction in the activation states of CBC-related enzymes (Fig. 4f). In comparison, in *2cpab* significantly higher activity was retained, with a negligible 5% loss of activation, suggesting that, under these conditions, 2-Cys Prx fully accounts for the CBC inactivation process (Fig. 4d and f). This observation, demonstrating the higher activation state of CBC following dark adaptation in *2cpab*, correlates with the reported maintenance of a highly reduced state of CBC enzymes in the dark (Doron et al. 2025). Intermediate inactivation values of 15%, 15% and 11% were recorded for *acht2*, *acht1/acht2*, and *acht1/acht2/acht4*, respectively (Fig. 4f), while no significant change in the deactivation state was observed for *acht1, acht4, acht1/acht4, trxl2.1* or *trxl2.2* (Fig. 4f, Supp. Fig. 8a). The fact that, *acht2-*containing lines did not fully mimic the *2cpab* phenotype suggests that additional limitations to carbon assimilation exist under low-light conditions, that are independent of ACHT2 but regulated by 2-Cys Prx.

When plants were reilluminated under moderate light intensity (140 µmol m⁻² s⁻¹), which closely approximates standard Arabidopsis growth conditions, the *2cpab* mutant exhibited a 22% decrease in the CBC activation state, and WT exhibited a 39% reduction (Fig. 4g), suggesting that oxidative deactivation is not solely regulated by 2-Cys Prx under these conditions. Mutants with an *acht2*-deficient background showed a CBC deactivation profile that was distinct from that of all other atypical Trx genotypes. More specifically, all *acht2*-containing lines (*acht2*, *acht1/acht2* and *acht1/acht2/acht4*) showed reduced CBC deactivation relative to WT and similar to *2cpab* (18%, 18%, and 16%, respectively). On the other hand, in *acht1*, *acht4* and *acht1*/*acht4*, CBC deactivation degrees were indistinguishable from the WT (Fig. 4g). Similarly, no significant changes in deactivation were noted in *trxl2* mutants (Fig. 4g and Supp. Fig. 8b).

To enable direct correlation of the redox kinetics of the electron transport chain with the degree of CBC activation, Fd oxidation was monitored following 1- and 6-min dark intervals, mirroring the conditions of the CBC inactivation assays. Notably, after 6 min of darkness, the *acht2* mutant maintained significantly accelerated Fd oxidation kinetics compared to the WT. Consequently, the increased availability of the electron acceptor pool was propagated upstream, resulting in markedly faster oxidation kinetics for both Pc and P700 (Supp. Fig. 9). These results flag ACHT2 as the primary mediator of oxidative inhibition of CBC enzymes.

## Discussion

In chloroplasts, both reductive and oxidative signals originate from the PETC; the reductive pathway is initiated by reduced Fd, whereas oxidative pathways depend on H₂O₂ production driven by the water-water cycle (Hipsch et al., 2026). Effective integration of thiol redox changes into a signaling module emerges from the coordinated interplay between reductive and counteracting oxidative activities, with both processes acting simultaneously on distinct subsets of target proteins to maintain a stable regulatory network (Danon et al., 2002; Doron et al., 2025). A central question in redox signaling is how specificity is achieved in a network composed of multiple proteins that are all exposed to opposing signals (i.e., reduced Fd and H₂O₂). It is assumed that the large number of chloroplastic Trxs, differing in midpoint reduction potential (Eₘ), binding affinities and effective concentrations (Schürmann et al., 2008; Dangoor et al., 2009), explains, at least in part, signaling specificity (Yoshida et al., 2015; Yoshida et al., 2018; Ojeda et al., 2018). However, the extent to which different Trxs contribute to signaling specificity rather than functional redundancy remains unclear. For example, examination of Trxs-target specificity revealed that distinct Trx isoforms exhibit high target specificity in vitro, despite significant functional redundancy *in vivo* (Yoshida et al., 2023). While Trx-f is uniquely efficient at reducing FBPase in biochemical assays, the near-normal growth of *trxf1/f2* mutants suggests that other Trx subtypes provide robust compensation at the whole-plant level (Yoshida et al., 2023). This work examined specificity versus functional redundancy among high-midpoint-potential Trxs implicated in oxidative signal transmission by analyzing electron-transfer and CO_2_ assimilation responses to dynamic light conditions in a set of CRISPR-generated mutant lines.

Given that redox modulation of enzymes involved in CO_2_ assimilation is reflected in limitations of the electron transport chain, physiological approaches to investigate the functional roles of various Trxs were adopted, integrating continuous in vivo physiological assessments that link these molecular states to their direct functional context. This avenue revealed that the ACHT/TRXL2 families do not operate as a redundant collective; instead, individual members possess distinct roles in governing chloroplast oxidative regulation. Specifically, ACHT2 was identified as a major determinant of carbon assimilation pathway activation during dark-light transitions. This role was reflected in the dynamics of Y(NA), Fd redox state, chlorophyll fluorescence and CO_2_ assimilation during light transitions (Fig. 2, 3 and 4) in *acht2*-containing mutants, which most closely resembled those of the *2cpab* mutant, in which the oxidative pathway is inhibited (Fig. 4). Notably, the lower Y(NA) observed in *acht2*-containing mutants, reflecting relief of the acceptor-side bottleneck, closely resembles the phenotype reported in plants overexpressing NTRC (Nikkanen et al., 2016), in which maintenance of 2-Cys Prx in a reduced state effectively blocks the oxidative pathway.

In an effort to capture target specificity, Yokochi et al. (2021) examined the oxidative activity of several Trxs by monitoring the redox states of key photosynthetic enzymes during light-to-dark transitions in several Trx mutant lines. Their analysis showed that TrxL2.1 oxidizes the γ-subunit of ATP synthase (CF₁-γ), whereas ACHT1 and/or ACHT2 contribute to the oxidation of FBPase. This functional specificity was further supported by ACHT2 over-expressing lines which demonstrated lower reduction levels (Fukushi et al., 2024). The current finding that ACHT2 plays a dominant role in CBC inactivation is consistent with its proposed role in FBPase oxidation, as FBPase activity is a key determinant of carbon assimilation rates (Tamoi et al., 2005; Tamoi et al., 2006; Rojas-González et al., 2015). However, because carbon assimilation is regulated by multiple enzymes and processes, some of which exert significant control over the Calvin cycle (Raines et al., 2003), modulation of FBPase activity alone is unlikely to fully account for the observed phenotypes.

No significant differences were observed in the measured photosynthetic parameters between WT and *trxl2* mutant lines, suggesting that although the TrxL2/2-Cys Prx redox cascade is involved in transmitting oxidative signals to chloroplast proteins (Yoshida et al., 2018), its activity does not govern the physiological regulation of the CBC during dark-to-light transitions. Given that dark-induced oxidation of the chloroplast ATP synthase CF₁-γ subunit is primarily mediated by TrxL2.1 (Yokochi et al., 2021), the WT-like phenotype of *trxl2* mutant lines suggests that ATP synthase reduction is unlikely to be the rate-limiting step for either CBC activation or the relief of acceptor-side limitation. Consistent with this, the CF1-γ subunit has been shown to rapidly reach a fully reduced state upon illumination, even under limited light conditions (Yoshida et al., 2014). Notably, *trxl2* mutant lines exhibited a WT-like pmf phenotype, characterized by a transient buildup of pmf during the light transition; this transient increase was attenuated in *acht2* mutant lines (Fig. 2g). These findings suggest that the transient accumulation of pmf during light transitions is driven primarily by limitations on CBC activity and subsequent ADP limitation, rather than by changes in proton conductivity across the thylakoid membrane.

The presented analysis suggested that ACHT4 exerts specific control over thylakoid electron transport dynamics. This was exemplified by the uniquely lower Y(ND) in the *acht4* mutant compared with the other lines, while similar Y(NA) values were maintained (Fig. 2f and Supp. Fig. 3). Crucially, the reduction in Y(ND) occurred despite the maintenance of a WT-like pmf in the *acht4* mutant (Fig. 2g), indicating that this regulation is uncoupled from the ΔpH-dependent photosynthetic control mechanism. Given that activation of photosynthetic control involves redox regulation via oxidation of specific thiols at the cytochrome b6f complex (Degen et al., 2024), these findings suggest that ACHT4 mediates redox signaling to the cytochrome b6f complex, a possibility warranting further investigation.

Plants lacking 2-Cys Prxs exhibit reduced growth and lower chlorophyll content compared with WT plants, suggesting an overall reduction in plant fitness (Awad et al., 2015; Ojeda et al., 2018). However, because 2-Cys Prx has been implicated in multiple functions, including antioxidant activity (Awad et al., 2015), chaperone activity (König et al., 2013) and thiol oxidase activity mediating the inactivation of photosynthetic enzymes (Pérez-Ruiz et al., 2017; Ojeda et al., 2018; Vaseghi et al., 2018; Yoshida et al., 2018), it remains unclear which of these functions is primarily responsible for the growth retardation observed in the mutant lines. The near-complete phenocopy of the *2cpab* defect in dark-induced CBC inactivation by *acht2* mutant lines suggests that loss of CBC activation/inactivation cycles during light-dark transitions has little impact on plant fitness under the tested conditions. The growth retardation observed in the *acht1/acht4* double mutant raises the possibility that the growth defect of *2cpab* plants is linked to the loss of specific 2-Cys Prx-dependent thiol oxidase activities mediated by ACHT1 and ACHT4. Given the established roles of ACHT4 in starch synthesis and ACHT1 in regulation of electron transport (Dangoor et al., 2012; Eliyahu et al., 2015), the severe *2cpab* phenotype may result from the simultaneous disruption of the regulation of these pathways rather than from the inhibition of dark inactivation of CBC activity.

In conclusion, physiological characterization of photosynthetic responses to dynamic light conditions in lines bearing mutant Trxs associated with the oxidative pathway, led to the identification of ACHT2 as a key regulator of CBC inactivation cycles. These findings provide a framework for exploring the ecological significance of redox regulation in plant adaptation to fluctuating light environments.

## Materials and Methods

### Plant materials and growth condition

Wild-type *Arabidopsis thaliana* (ecotype Col-0) was used in this study. Seeds were sown (Kultursubstrat, Klasmann-Deilmann, Geeste, Germany) and stratified at 4 °C, in the dark, for 48 h, to synchronize germination. Plants were subsequently cultivated in a controlled environmental chamber under long-day conditions (16-h light/8-h dark) with a photosynthetic photon flux density of 120 µmol photons m⁻² s⁻¹. Chamber parameters were maintained at 21 °C, 70% relative humidity, and ambient CO₂. All physiological measurements were performed on 3-4-week-old plants in a FytoScope FS-RI 1600 growth chamber (Photon Systems Instruments, Drásov, Czech Republic).

### CRISPR/Cas9-mediated genome editing

To generate the targeted knockout mutant lines, CRISPR-Cas9 binary vectors were assembled using the MoClo system (Weber et al., 2011, Engler et al., 2014). The binary vector cassette contained a functional Cas9 under a constitutive promoter (CaMV 35S), along with two, four or six gRNAs, kanamycin resistance and an mCherry fluorescent selection marker, which is selectively expressed in the seed coat (Omary et al., 2022). The final vectors were introduced into *Agrobacterium tumefaciens* strain GV3101 via heat shock, and T0 transgenic *Arabidopsis thaliana* plants were generated using the standard floral dip method (Clough and Bent, 1998). T1 transformants were initially identified by selecting seeds exhibiting red fluorescence under a fluorescence stereomicroscope. These T1 plants were cultivated, and the presence of targeted indels was confirmed via amplicon sequencing. Edited T1 individuals were then allowed to self-pollinate. In the subsequent T2 generation, seeds were screened by selecting non-red-fluorescent seeds indicating successful Mendelian segregation and complete loss of the Cas9 construct. These transgene-free T2 plants were genotyped to identify individuals strictly homozygous for the targeted mutations. CRISPR/Cas9-generated mutations were genotyped by PCR amplification of the target region in DNA extracted from plant leaves. Primers were designed to bind 120-400 bp away from the gRNAs. PCR products were analyzed by gel electrophoresis, followed by purification and targeted amplicon sequencing using either Sanger sequencing or Oxford Nanopore sequencing. Nanopore reads were aligned and analyzed using minimap2 (Li et al., 2018) and bcftools (Danecek et al., 2021) for variant caller. All physiological experiments were performed utilizing these verified, Cas9-free homozygous lines from the T2 and T3 generations.

### Measurements of plastocyanin, P700 and ferredoxin redox changes

The deconvoluted Pc, P700, and Fd redox changes were determined on detached leaves by NIR spectroscopy using the DUAL-KLAS-NIR spectrophotometer (Heinz Walz GmbH, Effeltrich, Germany, Klughammer & Schreiber, 2016). The NIRMAX script (Klughammer & Schreiber, 2016) was used to determine the maximal oxidation of Pc and P700, and the maximal reduction of Fd, for each tested leaf; maximal redox changes were used to normalize the data. Plants were dark-adapted for at least 1 hour, followed by illumination at 40 or 200 μmol photons m⁻² s⁻¹.

### Saturating pulse analysis of chlorophyll fluorescence and P700+

Quantum yields of PSI, Y(ND) and Y(NA), were determined upon stepwise increases in actinic light intensity, via saturating pulse analysis of the P700⁺ signal using the DUAL-KLAS-NIR spectrophotometer. Chlorophyll fluorescence was recorded simultaneously to calculate the quantum yield of PSII, NPQ and qL. To accurately capture the rapid photosynthetic transitions induced by illumination changes, a high-resolution pulse protocol was employed. Saturating pulses (7,800 μmol photons m⁻² s⁻¹) were applied at the onset of each new actinic light intensity, followed by three initial saturating pulses at 5-s intervals, and subsequent pulses every 15 s for the remaining light period.

### Electrochromic shift (ECS) measurements

Changes in the electrochromic shift (ECS) signal were monitored utilizing a DUAL-KLAS spectrophotometer equipped with a P515/535 accessory module (Schreiber & Klughammer, 2008). Consistent with the redox kinetics assays, plants were dark-adapted for at least 1 h. To evaluate the pmf and the proton conductivity of the chloroplast ATP synthase (gH+), dark interval relaxation kinetics (DIRK) was recorded during the actinic light curve. To precisely align the ECS data with the redox analyses, the timing of the DIRK measurements perfectly aligned with the timing of the saturating pulse experiments. The test protocol used rapid dark intervals of 300 ms and a measuring light pulsed at 2,000 Hz. ΔECS_T_ values were normalized to the amplitude measured upon a 20 µs single-turnover saturating flash applied to the dark-adapted leaves prior to the actinic light curve (Mathiot & Alric, 2021). The total pmf was estimated from the ΔECS_T_ amplitude, calculated as the difference between the steady-state ECS signal in the light and the y0 baseline value derived from a first-order exponential fit of the decay kinetics. gH+ was subsequently determined as the inverse of the exponential decay time constant. All data processing and first-order exponential curve fitting were performed using a custom Python script in PyCharm.

### Gas exchange measurements

Carbon assimilation rates were measured utilizing a LI-6800 portable photosynthesis system equipped with a 6800-17 Small Plant Chamber (LI-COR Biosciences, Lincoln, NE, USA). Environmental conditions within the measurement cuvette were strictly maintained at a temperature of 21 °C, a reference CO₂ concentration of 420 ppm, a relative humidity of 70%, and a constant airflow rate of 700 µmol s⁻¹. To ensure optimal gas mixing, the chamber fan speed was set to 5,000 rpm. Furthermore, the instruments’ infrared gas analyzers (IRGAs) were recalibrated using the automatic matching function immediately before each plant measurement.

Measurements were performed on three intact plants (3-4 weeks old, pre-flowering stage) grown in 65-mm pots (LI-COR part no. 610-09646) specifically designed to interface with the chamber. Gas-exchange variables were continuously recorded at 5-s intervals throughout each experimental run. To accurately quantify assimilation on a per-leaf-area basis, total rosette area was determined post-measurement. Plants were imaged utilizing the CropReporter (PhenoVation B.V., Wageningen, The Netherlands), and the total leaf area was extracted via image analysis in MATLAB. Absolute assimilation rates were subsequently normalized to the calculated area.

### Quantification of Calvin-Benson cycle (CBC) activation

Photosynthesis induction kinetics were recorded during a sequential light-dark transition protocol. To establish a proper baseline, plants were first acclimated to the specific light intensity tested and then continuously monitored throughout the sequential dark intervals. Actinic illumination was provided at an intensity of either 40 or 140 µmol photons m⁻² s⁻¹. To evaluate the kinetics of carbon assimilation recovery, the continuous light phase was interrupted by two distinct dark adaptation intervals: an initial 1-min dark period, followed by a subsequent 6-min dark period. Plants were re-illuminated to steady-state after each dark interval. To ensure unbiased quantification of induction kinetics, initial slopes of carbon assimilation were determined using a custom Python script. For each dark-to-light transition, the script iteratively identified the continuous linear region that yielded the maximal R^2^, with the search initiated from the onset of illumination (minimum of n=7 points). To account for inherent variations in absolute photosynthetic capacity between plant lines, each induction slope was normalized to the steady-state plateau reached during that specific 4-min illumination phase. The plateau was defined as the terminal segment of the light period (minimum n=10 points) exhibiting the minimum variance, ensuring a stable and representative baseline for each cycle. The relative activation degree of the CBC was then calculated as the ratio of the normalized induction slope following the 6-min dark period to the normalized slope following the 1-min dark period, expressed as a percentage:

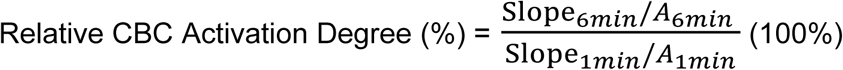

### Statistical analysis and data visualization

Statistical analyses and data visualization were performed using R (version RStudio 2024.12.1) within the RStudio environment, JASP (Version 0.95.4.0, https://jasp-stats.org/) and custom MATLAB scripts. To evaluate the significance of observed differences across genotypes, a one-way ANOVA was employed, followed by Tukey-Kramer post-hoc tests for multiple comparisons. Statistical significance was established at p < 0.05. For independent pairwise comparisons of mutant lines against the wild type (Fig. 2, Supp. Fig. 2 and 3), Welch’s *t*-test with a Bonferroni correction was applied. Significance for these comparisons is marked as p < 0.05 (*), < 0.01 (**), < 0.001 (***).

For all box plots, the box encompasses the interquartile range (IQR) from the 25th to the 75th percentile, with the horizontal line representing the median and the red dot indicating the mean value. Whiskers illustrate the distribution of data extending beyond the interquartile range. To ensure full data transparency, all individual biological replicates were overlaid as semi-transparent grey circles. Genotypic groups categorized by the same letter are not significantly different according to the post-hoc analysis. To ensure data robustness, outliers were identified and removed independently for each genotype using the standard IQR method. Outlier data points were excluded from the final datasets. To verify that this filtering process did not artificially skew the conclusions, all downstream statistical analyses were also independently performed on the raw, unadjusted datasets. Both the IQR-filtered and raw datasets yielded identical statistical significance outcomes, confirming that the removal of outliers did not alter the significance of the results.

## Supporting information

Supplementary figures steinberg et al., 2026

## Acknowledgments

The authors thank Prof. Idan Efroni, who kindly provided the indicated MoClo-compatible Cas9 and sgRNA backbone plasmids

