## Supplementary figures steinberg et al., 2026 for "ACHT2 deactivates carbon assimilation during light-dark transition"

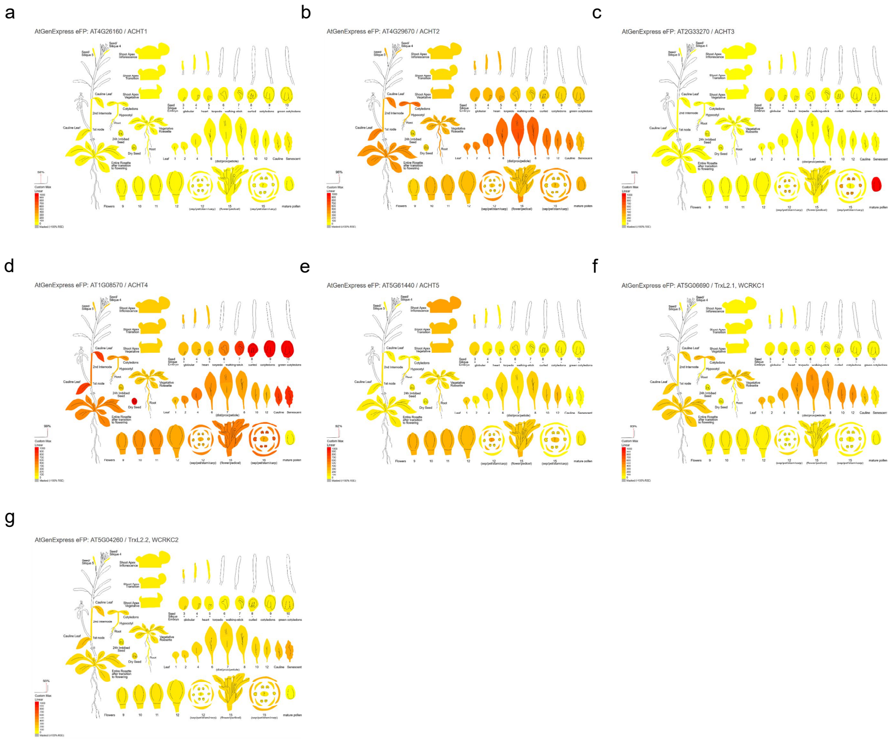


**Supplementary Figure 1. Tissue specific expression profiles of atypical thioredoxin genes.** Panels display the expression levels of **(a)** *acht1*, **(b)** *acht2*, **(c)** *acht3*, **(d)** *acht4*, **(e)** *acht5*, **(f)** *trxl2.1*, and **(g)** *trxl2.2* mapped across vegetative and reproductive tissues. To enable direct visual comparison of transcript accumulation, expression intensity was scaled to a uniform custom color gradient across all panels, ranging from 0 to a maximum of 1000. Transcript abundance was visualized using the Arabidopsis eFP viewer (https://bar.utoronto.ca/eplant/).

| Gene | INDEL | Position |
|---|---|---|
| *acht1_13.2* | -7 | Exon 1 +306 |
| *acht1_18.2* | +1 | Exon 1 +306 |
| *acht2_1.4* | +1 | Exon 1 + 235 |
| *acht2_13.2* | +1 | Exon 1 +235 |
| *acht2_13.7* | -54 | Exon 1-2 +234 |
| *acht4_15.1* | -127 | Exon 2-3 +410 |
| *acht4_16.6* | +1 | Exon 1 +410 |
| *acht1* + | +1 | Exon 1 +306 |
| *acht4* | +1 | Exon 1 +410 |
| *acht1* + | +1 | Exon 1 +306 |
| *acht2* | -54 | Exon 1-2 +234 |
| *acht1* + | +1 | Exon 1 +306 |
| *acht2* + | -53 | Exon 1-2 +233 |
| *acht4* | -127 | Exon 2-3 +410 |
| *trxl2.1* | +8 | Exon 1 +289 |
| *trxl2.2_9.1* | -123 | Exon 1 +146 |
| *trxl2.2_16.1* | +1 | Exon 1 +268 |

**Supplementary Table 1. Summary of CRISPR/Cas9-generated mutations in *ACHT* and *Trx-like* genes.** Columns indicate the target gene, the nature of the specific insertion/deletion (indel), and the intragenic position of the mutation.

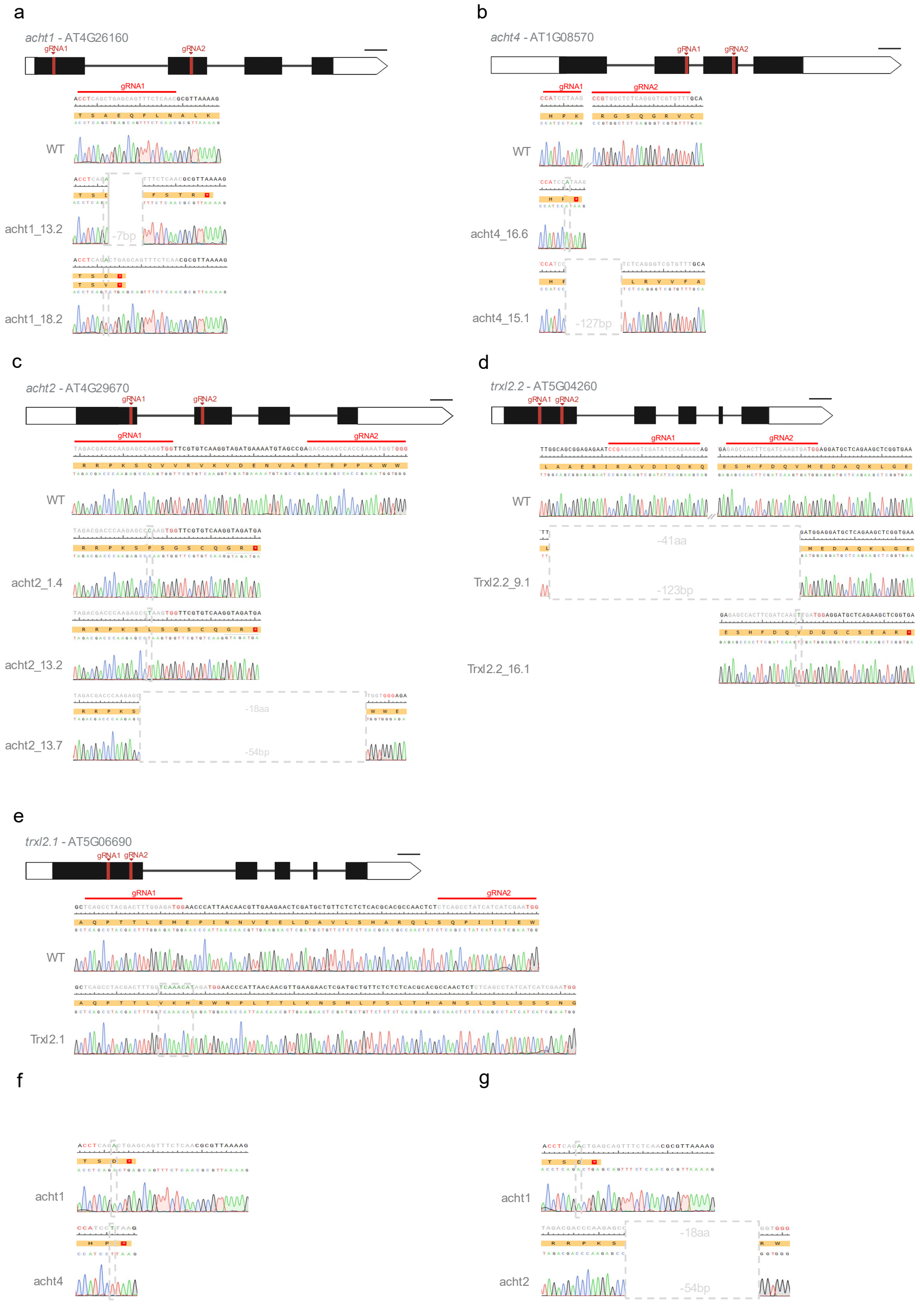


**Supplementary Figure 2. Sanger sequencing validation of targeted mutations in *ACHT* and *Trx-like* genes.** Chromatograms display the specific genetic lesions for mutant lines: **(a)** *acht1*, **(b)** *acht4*, **(c)** *acht2*, **(d)** *trxl2.2*, **(e)** *trxl2.1*, as well as the double mutant lines **(f)** *acht1*/*acht4* and **(g)** *acht1*/*acht2*. For each panel, WT sequence is aligned with the mutant chromatogram for comparison. Cas9 target sites are indicated in red on the respective gene structures. Within the sequence alignments, nucleotides are color-coded as follows: gray denotes the sgRNA sequence, red indicates the PAM, and green highlights inserted nucleotides. All resulting indels are enclosed within dashed gray rectangles.

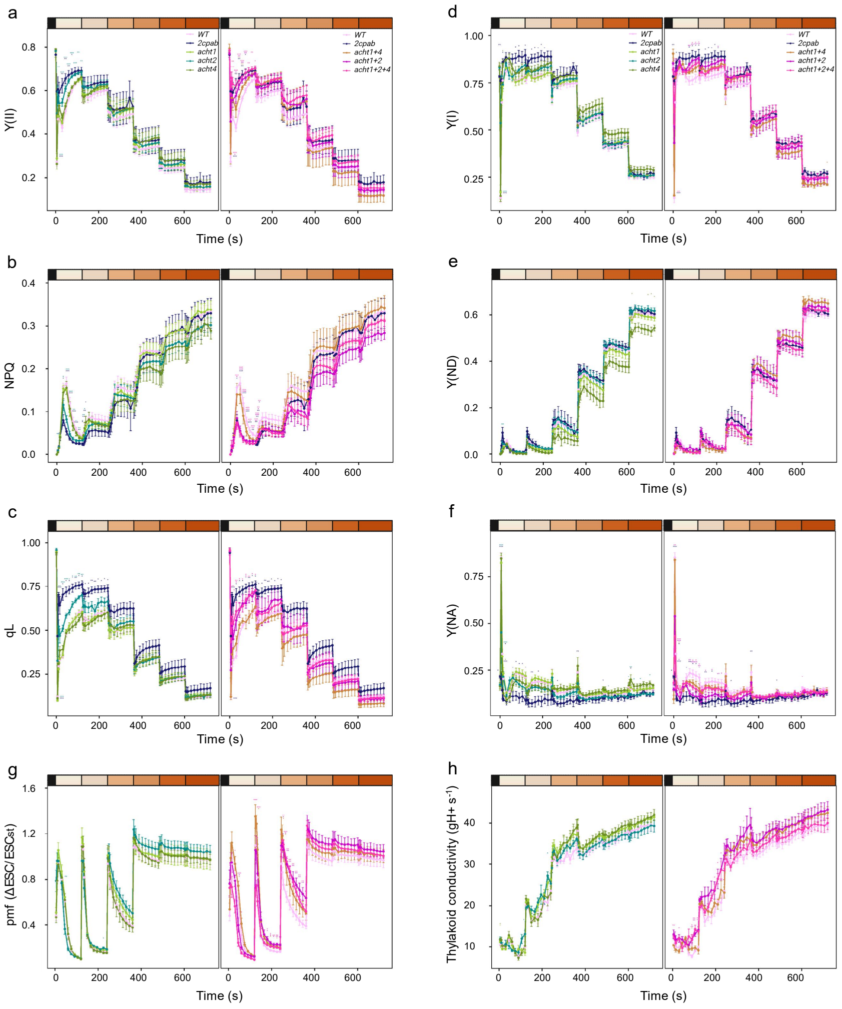


**Supplementary Figure 3. Comprehensive rapid light response curves of photosynthetic parameters across the *acht* mutant lines .** Analysis of kinetic photosynthetic parameters under a complete gradient of actinic light intensities (0, 35, 80, 160, 350, 650, and 1200 µmol photons $m^{-2}$ $s^{-1}$). Panels display the full induction response for **(a)** quantum yield of PSII (Y(II)), **(b)** non-photochemical quenching (NPQ), **(c)** fraction of open PSII reaction centers (qL), **(d)** quantum yield of PSI (Y(I)) , **(e)** PSI donor-side limitation (Y(ND)), **(f)** PSI acceptor-side limitation (Y(NA)), **(g)** proton motive force (pmf), and **(h)** thylakoid proton conductance (gH+)**.** Data represent the mean ± SE (n = 5-20 biological replicates). Statistical significance was assessed for panels (a-h) was assessed using Welch's *t*-test with Bonferroni correction for multiple comparisons against the wild type and is marked as p < 0.05 (*), p < 0.01 (**), p < 0.001 (***).

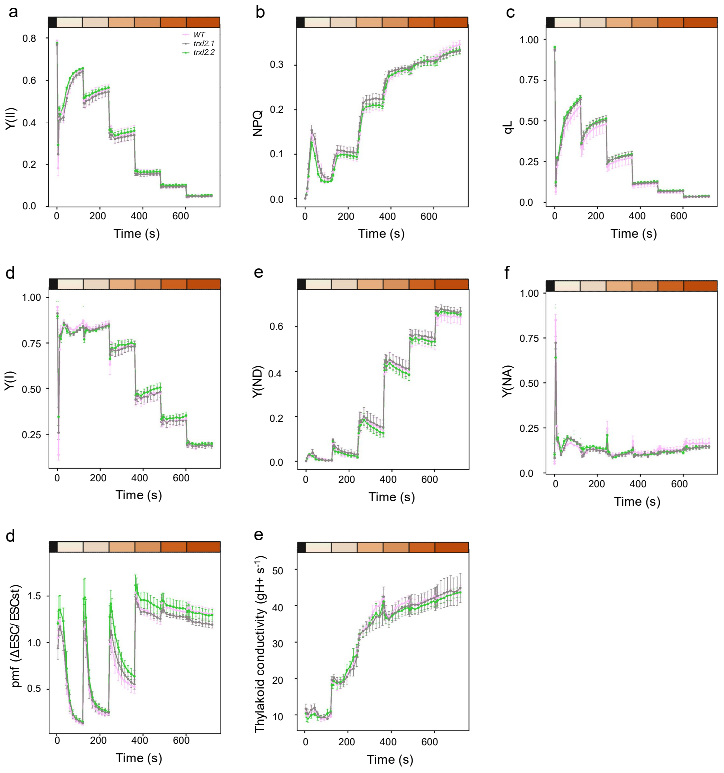


**Supplementary Figure 4. Comprehensive rapid light response curves of photosynthetic parameters across the *trxl2* mutant lines .** Analysis of kinetic photosynthetic parameters under a complete gradient of actinic light intensities (0, 35, 80, 160, 350, 650, and 1200 µmol photons $m^{-2}$ $s^{-1}$). **(a)** Panels display the full induction response for quantum yield of PSII (Y(II)), **(b)** Non-photochemical quenching (NPQ), **(c)** Fraction of open PSII reaction centers (qL), **(d)** Quantum yield of PSI (Y(I)), **(e)** PSI donor-side limitation (Y(ND)), **(f)** PSI acceptor-side limitation (Y(NA)), **(g)** Proton motive force (pmf), and **(h)** thylakoid proton conductance (gH+). Data represent the mean ± SE (n = 4-6 biological replicates). Statistical significance was assessed for panels (a-h) was assessed using Welch's *t*-test with Bonferroni correction for multiple comparisons against the wild type and is marked as $p < 0.05$ (*), $p < 0.01$ (**), $p < 0.001$ (***).

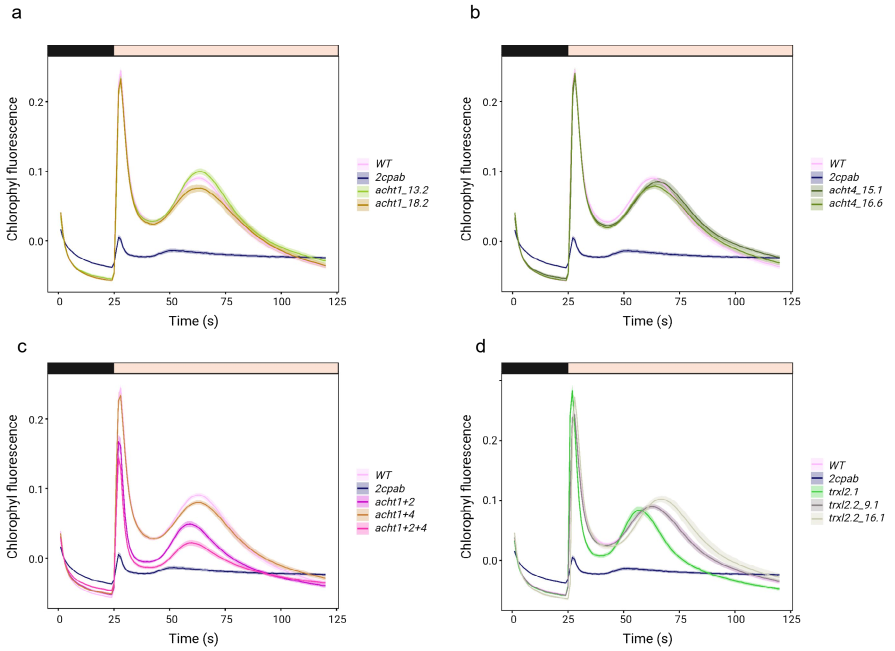


**Supplementary Figure 5. Chlorophyll fluorescence induction kinetics under dark to low light transition. (a-d)** Analysis of kinetic photosynthetic induction curve across the independent mutant lines under dark to low light (35 µmol photons $m^{-2} s^{-1}$). Data are presented as the mean ± SE (n = 8 - 32 biological replicates).

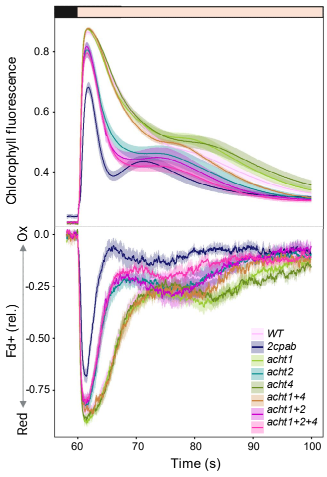


**Supplementary Figure 6. The activation state of Fd strongly correlates with chlorophyll fluorescence quenching.** Analysis of the relationship between the reduction state of the downstream electron acceptor, Fd, and chlorophyll fluorescence quenching dynamics. Data are presented as the mean ± SE (n = 8 - 32 biological replicates).

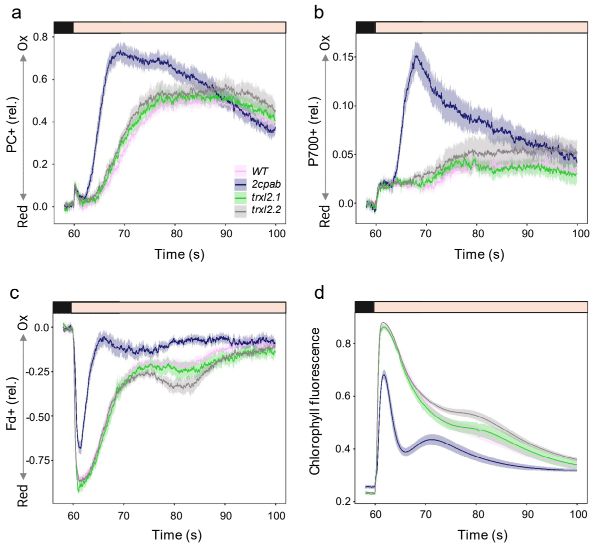


**Supplementary Figure 7. Chlorophyll fluorescence and redox state of Pc, p700 and Fd real-time induction kinetics under dark to low light transition.** Analysis of kinetic photosynthetic induction across the *trxl2* mutant lines under dark to low light (35 µmol photons $m^{-2}$ $s^{-1}$). **(a)** Fraction of oxidized Pc. **(b)** Fraction of oxidized p700. **(c)** Fraction of reduced Fd, **(d)** Chlorophyll fluorescence induction curve. Data are presented as the mean ± SE (n = 8 - 32 biological replicates).

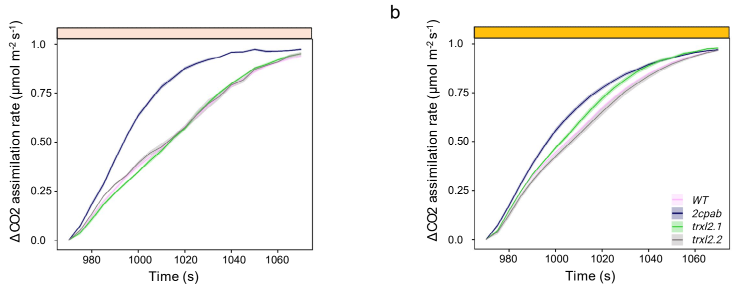


**Supplementary Figure 8. Initial induction rates of CO• assimilation after 6 min of dark adaptation.** Transient assimilation rates of *trxl2* were measured under subsequent **(a)** moderate or **(b)** low light intensities. To evaluate relative induction kinetics, all data were normalized to the final steady-state assimilation rate achieved under each respective light condition.

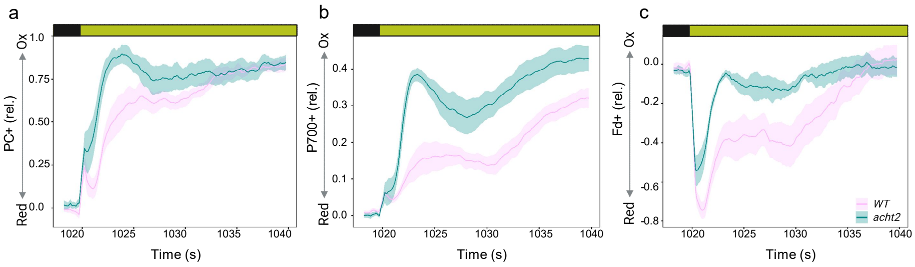


**Supplementary Figure 9. Real-time induction kinetics of PC, P700, and Fd redox states during dark-to-light transitions.** Following 6 min of dark adaptation, kinetic photosynthetic induction was recorded across the WT and *acht2* mutant lines upon transition to moderate light intensity (200 µmol photons $m^{-2}$ $s^{-1}$). Panels display the transient relative redox states of **(a)** the fraction of oxidized Pc, **(b)** the fraction of oxidized p700, and **(c)** the fraction of reduced Fd. Data are presented as the mean ± SE (n = 4 - 5 biological replicates).
